# Dynamic Human Gut Microbiome and Immune Shifts During an Immersive Psychosocial Therapeutic Program

**DOI:** 10.1101/2024.06.26.600881

**Authors:** Xin Zhou, Ariel B. Ganz, Andre Rayner, Tess Yan Cheng, Haley Oba, Benjamin Rolnik, Samuel Lancaster, Xinrui Lu, Yizhou Li, Jethro S. Johnson, Rebecca Hoyd, Daniel J. Spakowicz, George M. Slavich, Michael P. Snyder

## Abstract

**Background:** Depression is a leading cause of disability worldwide yet its underlying factors, particularly microbial associations, are poorly understood.

**Methods:** We examined the longitudinal interplay between the microbiome and immune system in the context of depression during an immersive psychosocial intervention. 142 multi-omics samples were collected from 52 well-characterized participants before, during, and three months after a nine-day inquiry-based stress reduction program.

**Results:** We found that depression was associated with both an increased presence of putatively pathogenic bacteria and reduced microbial beta-diversity. Following the intervention, we observed reductions in neuroinflammatory cytokines and improvements in several mental health indicators. Interestingly, participants with a *Prevotella*-dominant microbiome showed milder symptoms when depressed, along with a more resilient microbiome and more favorable inflammatory cytokine profile, including reduced levels of CXCL-1.

**Conclusions:** Our findings reveal a protective link between the Prevotella-dominant microbiome and depression, associated with a less inflammatory environment and moderated symptoms. These insights, coupled with observed improvements in neuroinflammatory markers and mental health from the intervention, highlight potential avenues for microbiome-targeted therapies in depression management.

## INTRODUCTION

Depression is a highly prevalent and economically burdensome psychiatric condition that is associated with a variety of physical health conditions^1^, and this situation has been exacerbated by the recent COVID-19 pandemic^2–6^. This condition affects a wide range of individuals across ages, genders, and geographical locations^7–9^. Despite its substantial societal impact, much remains unknown about the underlying biology of depression. This gap stems in part from the inherent molecular complexity of the human brain and behavior^10–12^, coupled with the challenges of replicating psychiatric disorders in animal models^13–15^, and the reliance on self-reported diagnostic tools like the Beck Depression Inventory-II (BDI-II)^16^ and the Patient Health Questionnaire (PHQ-9)^17^, which may introduces biases. Furthermore, our insights into depression have been constrained by the limited application of longitudinal multi-omics profiling studies, which are crucial for unraveling the complex molecular and cellular mechanisms underlying mental health disorders^18,19^. These collective challenges have hindered our understanding of the underlying molecular and cellular mechanisms of depression and have consequently impeded the advancement of novel pharmacological and psychotherapeutic interventions for major depressive disorder^20^.

Recent studies have highlighted a significant bi-directional association between depression and inflammation, shedding light on the frequent co-occurrence of depression with various inflammatory disorders^21,22^. This association is often viewed through frameworks such as the Social Signal Transduction Theory of Depression, which suggests that psychosocial stressors can trigger an inflammatory response, elevating depression risk in susceptible individuals^1^. The interaction between depression and inflammation is complex and reciprocal: inflammation can precipitate depressive symptoms^23,24^, and, in addition, depression can intensify inflammation through behavioral and physiological pathways^25,26^. This reciprocal relationship highlights the intricate connection between mental and physical health. Importantly, psychosocial interventions have emerged as effective in bolstering immune function, presenting a viable alternative to traditional antidepressants in managing depression-associated inflammation^27–29^.

Recognizing the dynamic between depression, inflammation, and immunity necessitates examining the gut microbiome’s impact on mental health. The gut microbiome, a key regulator of the immune system, significantly affects human behavior via the gut-brain axis^30–33^. Research indicates that psychiatric and behavioral disorders, including addiction^34–36^, depression^37–40^, aggression^41^, and impaired social cognition^42,43^ correlate with notable microbiome alterations. These changes are deeply intertwined with brain function and behavior by regulation of key metabolites such as serotonin (5-HT)^44,45^, tryptophan^35,46–48^, and γ-Aminobutyric acid (GABA)^49^, pivotal for neurotransmission, mood, cognition, and stress response. Serotonin, targeted by many antidepressants^50,51^, influences a wide range of psychological and physiological functions. Tryptophan is a precursor for serotonin synthesis, thus influencing serotonin levels and, consequently, mood and emotional states^52,53^. GABA, as the primary inhibitory neurotransmitter in the brain^54,55^, plays a key role in reducing neuronal excitability throughout the nervous system^56–59^, impacting processes like anxiety regulation and stress response. The microbiome extends its influence to the immune system, notably in cytokine regulation^48,60–62^, crucial for neuroinflammation and neural-immune interactions^63,64^. Furthermore, certain cytokines and chemokines, influenced by the microbiome, play a pivotal role in signal transduction processes^56,65–67^. These observations underscore the multifaceted impact of the gut microbiome on brain function and behavior, highlighting its role in both metabolic regulation and direct immune modulation.

Recent research has established the gut microbiome’s causal relationship with depression, as evidenced by experiments transferring microbiomes from depressed patients to mice, leading to depressive-like behaviors and altered metabolism^68–70^. Specifically, bacteria such as *Escherichia* have been linked to promoting depressive symptoms^69,71^. These effects may be mediated through the modulation of the hypothalamic-pituitary-adrenal (HPA) axis and cytokine production^71^, and significantly influence cytokine production, including Brain-Derived Neurotrophic Factor (BDNF)^72^ and Interleukin-6 (IL-6)^71^. Furthermore, the microbiome’s influence on the metabolism and efficacy of antidepressants has emerged as a significant area of interest, highlighting its potential to shape psychotherapeutic treatment outcomes^47,73–75^.

The longitudinal interplay between the microbiome and immune system in the context of depression, and in response to an immersive psychosocial intervention, remains unexplored, largely due to differences in human and animal immune and mental health systems^76^, and the impracticality of applying psychotherapeutic strategies such as self-inquiry and meditation in animal models. Research indicates that microbiome compositions, summarized by enterotypes like *Bacteroidetes*, *Firmicutes*, or *Prevotella*, are crucial for nutrient processing, inflammatory responses, and drug metabolism^77–81^. Specifically, *Firmicutes* ^82^ and *Prevotella*^83^ enterotypes have been linked to mental health, with the latter associated with increased positive emotions, though further research is required. To explore the interactions among the human microbiome, immune system, and depressive symptoms, we conducted a longitudinal study of participants going through a highly immersive, inquiry-based stress reduction program. This approach, integrating gut microbiome and plasma cytokine analyses with mental health assessments before and after the program, provides novel insights into microbiome-host dynamics during an intervention that has known therapeutic benefits.

## RESULTS

Fifty-nine individuals were recruited under Stanford IRB 48982, excluding those with cancer or steroid use. They attended a 9-day intensive inquiry-based stress reduction program at Ojai Valley Inn, California. Samples were collected upon arrival (T1), after the stay (T2), and one (T3) and three months (T4) post-retreat, totaling 142 stool and 123 plasma samples. (**Fig. 1A**) In addition to collecting biological samples, we conducted an in-depth assessment of participants’ mental health. This included depression, anxiety, perceived stress, and psychosocial indicators of well-being. The initial depression status of the participants was classified into two categories: "depressed" and "non-depressed." This categorization was based on their total BDI-II (Beck Depression Inventory-II) score, using a score of 14 and above as the threshold for being depressed, as per standard convention^16^.

**Figure 1.**
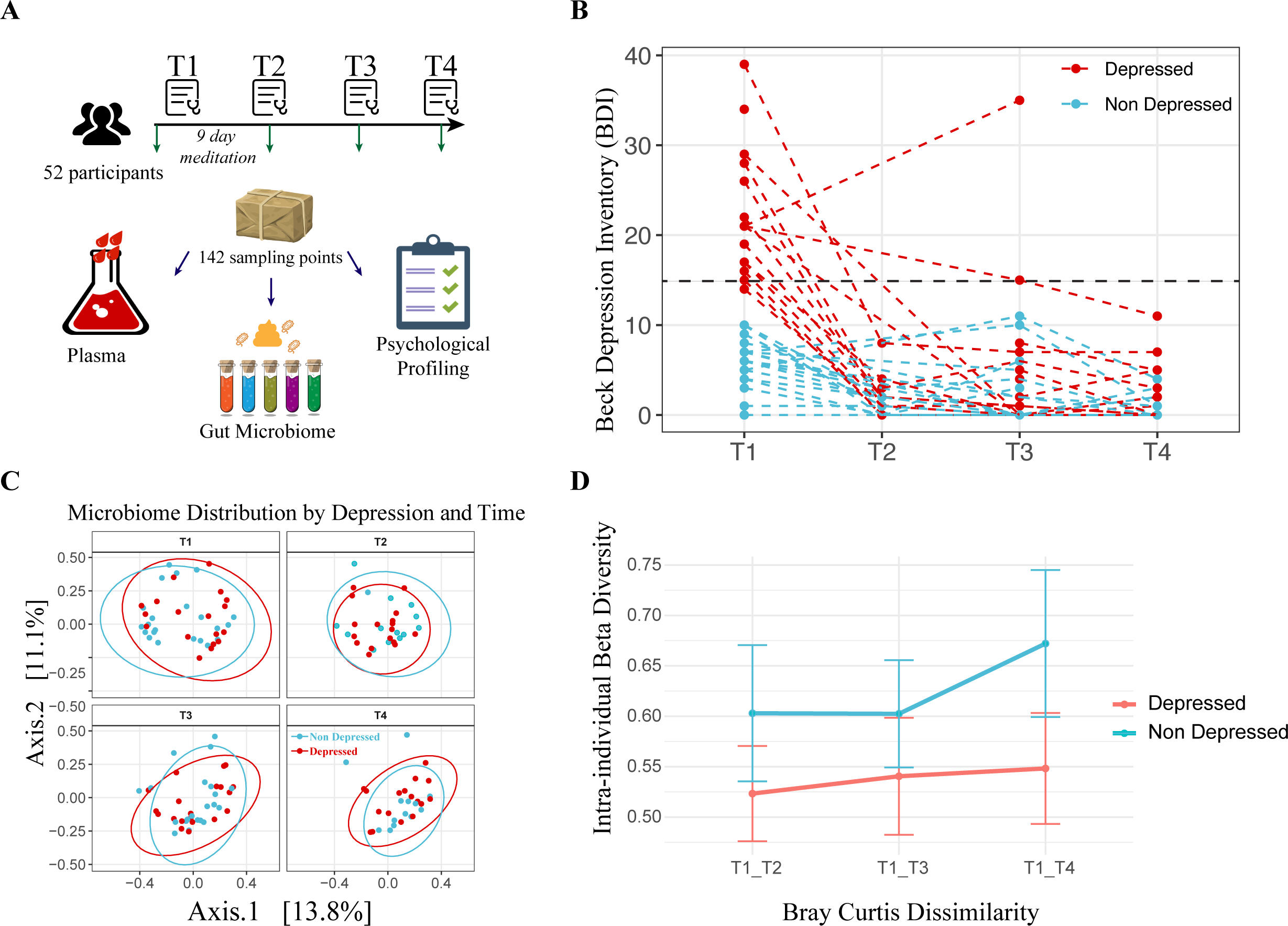
Study Design, Depression Trajectories, and Microbiome Composition Analysis. **(A)** Cartoon representation of the study’s methodology (Image adapted from *Pixabay*). **(B)** Beck Depression Inventory-II (BDI-II) scores of participants across four time points, with each dashed line corresponding to one individual. The color coding indicates whether the individual was depressed (red) or non-depressed (blue) at baseline. **(C)** Principal Coordinates Analysis (PCoA) of microbiome composition at the four time points. The first two axes are plotted, with the variance captured annotated along each axis. Red and blue colors denote the initial depression status of the participants, with red indicating depressed and blue indicating non-depressed individuals. **(D)** Intra-individual dissimilarity between T1 and the rest of the program.

Initially, 21 participants were identified as depressed based on BDI-II scores. Of these 21 participants, a full 20 of them (95.24%) exhibited a significant decrease in BDI-II scores post-program, with the reduction sustained over subsequent assessments. For the study’s duration, those classified as depressed at T1 remained in the "Depressed" group for analysis purposes, regardless of any changes in depression status at later time points (T2, T3, or T4). (**Fig. 1B**)

A Permutational Multivariate Analysis of Variance (PERMANOVA) revealed that the initial depression status accounted for a small yet statistically significant proportion of the variation in gut microbiome composition (R2 = 1.4%, Pr (>F) = 0.004). Yet, this did not significantly overshadow intrapersonal variations (2.3% variance, p = 0.21), which were not consistent across participants (**Fig. 1C, S1A**). For the non-depressed group, participation in the program (i.e., meditation) was associated with a substantial increase in microbial richness. This effect was measured using the Chao1 index, which showed a rise from the start to the end of the program (T1-T3: beta=35.06, p = 0.099), as well as one month later (T1-T4: beta = 84.84, p = 0.0016). Additionally, we observed a trend toward increased intraindividual beta-diversity (p=0.078) (**Fig. 1D**), which is consistent with findings from the largest meta-analysis on the subject^38^. This finding suggests that the immersive intervention program contributed to a mild, and potentially beneficial, increase in microbial diversity^40^, a change not observed for those in the depressed subgroup. (**Fig. S1B**)

To specifically address the longitudinal design and zero-inflated nature of our microbiome data, a separate analytical strategy was employed using a two-part Zero-Inflated Beta Regression model with Random effects (ZIBR)^84^. This approach identified three genera—*Solobacterium*, *Anaerofilum*, and *Escherichia/Shigella*—that exhibited differential distribution based on depression status across time (**Table S1**). Notably, while *Solobacterium* has previously been reported to increase among academic-related chronic stress among young students^85^, our data reveal a significant increase in the gut microbiome of depressed individuals. Such findings suggest translocation of pathogens from one body site to another during disease stage. *Escherichia/Shigella*^37,86–88^ and *Anaerofilum*^86,89^ have also been previously associated with depression and stress, thus providing an external validation for the model. This finding is consistent with our broader understanding that the gut microbiome’s association with mental depression appears to be characterized by the small to modest increase in pathogens, which, although impactful, represent only a minor fraction of the total microbiome, rather than systematic shifts in the core community.

Although our differential abundance analysis did not reveal significant pathogen overgrowth in individuals with depressive symptoms, we investigated whether enterotypes—distinct microbial configurations defined by specific bacterial genera dominance^90^—were related to changes in depression status over time. Enterotypes are known to significantly influence nutrient metabolism^91–93^, the immune environment^81,94–96^, and disease onset and treatment^97–100^, suggesting their potential role in depression dynamics. Combined with prior knowledge^90,101^, we grouped our cohort into high *Bacteroides*, high *Prevotella* and a *Firmicutes* enriched cluster that are low for the formal mentioned two genera but high for *Ruminococcus*. (**Fig. 2A, S2A**), leading to the classification into *Bacteroides* (Ba), *Ruminococcus* (Ru), and *Prevotella* (Pr) enterotypes. Our results show that these enterotypes remained stable throughout the study, indicating that short-term interventions like meditation may not significantly alter these established microbial communities. (**Fig. S2B**)

**Figure 2.**
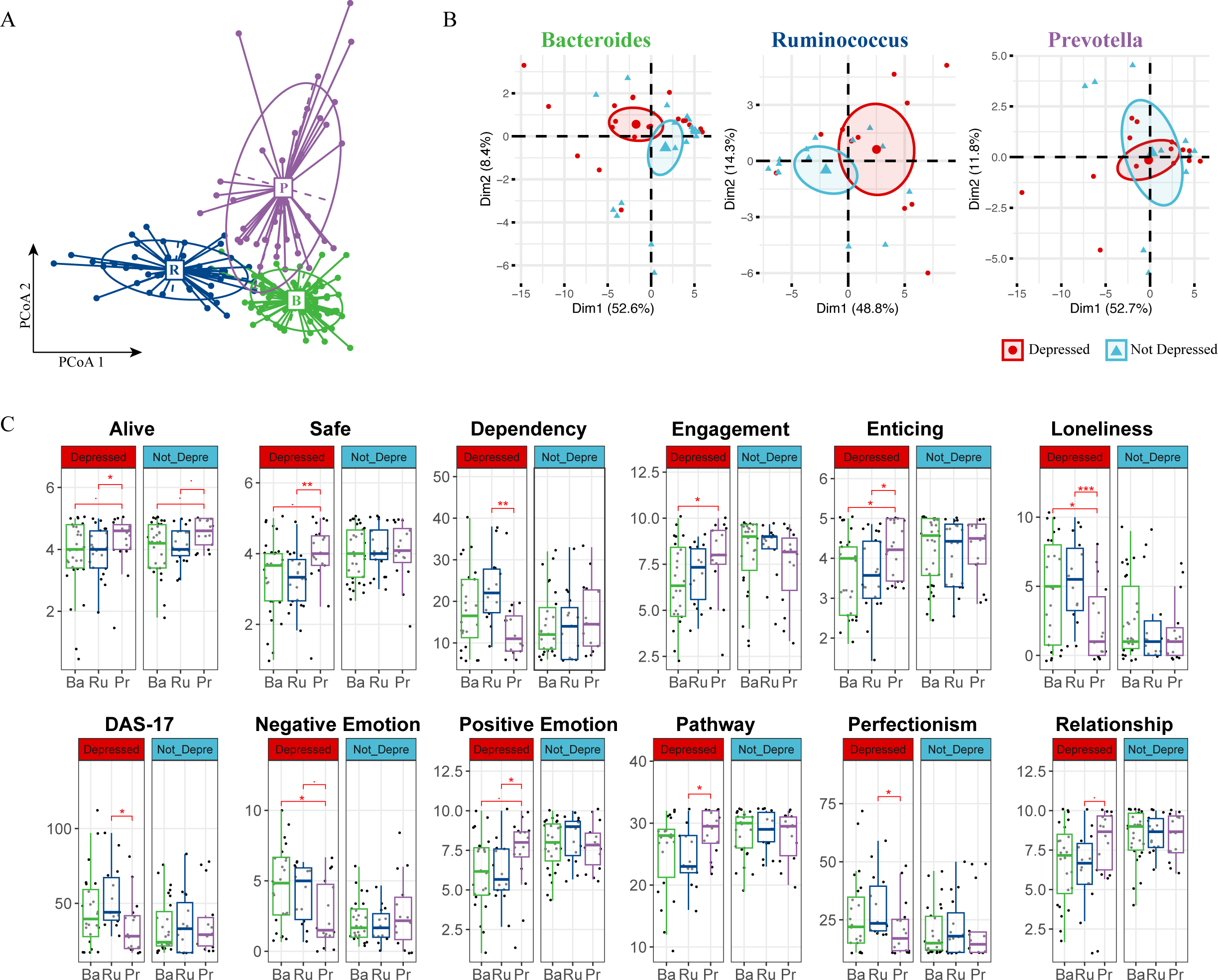
Enterotype Analysis Reveals Potential Beneficial Role of Prevotella in Depression. **(A)** Enterotype Classification of Microbiome Samples. Samples were clustered into three enterotypes. Each cluster is represented by a predominant genus, detailed in Supplementary Figure S2a. Colors represent different enterotypes: *Ruminococcus* (R, blue), *Bacteroidetes* (B, green), and *Prevotella* (P, purple). **(B)** PCA of Psychometric Parameters Based on Depression Status. Samples are colored based on each individual’s initial depression status (red for depressed, blue for non-depressed). The first two principal components (PCs) are plotted, with the variance explained by each PC annotated. **(C)** Pairwise Comparison of Psychometric Data by Enterotype and Depression Status. Comparison across *Ruminococcus* (Ru, blue), *Bacteroidetes* (Ba, green), and *Prevotella* (Pe, purple) enterotypes. A two-sided Student’s T-test was used for each comparison. Significance levels are indicated as follows: p < 0.1 (.), p < 0.05 (*), p < 0.01 (**), p < 0.005 (***).

Our psychometric data analysis revealed marked differences between depressed and non-depressed participants within the *Bacteroides* and *Ruminococcus* enterotypes, as shown by distinct clustering in Principal Component Analysis (PCA); such differentiation was absent in the *Prevotella* group. (**Fig. 2B**). The similar variance explained by the first two principal components suggests that individuals within the *Prevotella* enterotype, regardless of depression status, exhibit comparable psychometric profiles. Further analysis revealed a unique trend within the *Prevotella* group: individuals scored higher on feeling "alive," a pattern maintained across depression statuses. **(Fig. 2C).** Additionally, depressed individuals within the *Prevotella* enterotype reported a greater sense of safety, enticement, and positive emotions. They also noted higher levels of engagement and relationship satisfaction, coupled with lower tendencies towards dependency, loneliness, and perfectionism. Their scores on the Dysfunctional Attitudes Scale (DAS17) were also consistently lower. **(Fig. 2C).** The observations specific to the depressed individuals within the *Prevotella* group are not due to an overrepresentation of *Prevotella* in either the depressed or non-depressed groups **(Fig. S2B)**, nor did we find a statistically different average BDI-II score at the beginning of the program or throughout its entirety for the *Prevotella* group. In addition, we did not identify any significant association of enterotype with participants’ Adverse Childhood Experiences (ACEs) score (**Fig. S2C**); in fact, none of the above-mentioned psychometric parameters were hierarchically clustered with ACEs. (**Fig. S2D**)

Acknowledging the positive psychometric outcomes linked to the Prevotella enterotype, we sought to investigate the potential associations involving enterotypes, baseline immune responses, and their influence on depressive symptoms. To understand the immune profile associated with depression further, we examined cytokines, chemokines, and growth factors in date-matched plasma samples using an 80-plex Luminex assay. Using a PERMANOVA test that analyzed variance in cytokine data by timepoints, enterotype, and depression status across 9,999 permutations, we identified a significant association between systemic inflammation, as reflected in cytokine levels, and depression status (R^2^ = 1.91%, Pr > F = 0.005). Intriguingly, enterotype also contributed to a minor, yet statistically significant, variation in cytokine levels (R^2^ = 3.84%, Pr > F = 0.001).

Given the observed variance in cytokine levels influenced by both enterotype and depression status, we undertook hierarchical clustering of the cytokine data, focusing on the average values across different enterotypes and depression states. This was specifically to assess whether the Prevotella enterotype in depressed individuals exhibits a cytokine profile similar to that of non-depressed groups when considering all cytokines systematically. Indeed, our analysis revealed that depressed individuals with the Prevotella enterotype indeed clustered with three enterotypes from the non-depressed group, indicating a similar overall inflammatory profile (**Fig.3A**).

**Figure 3.**
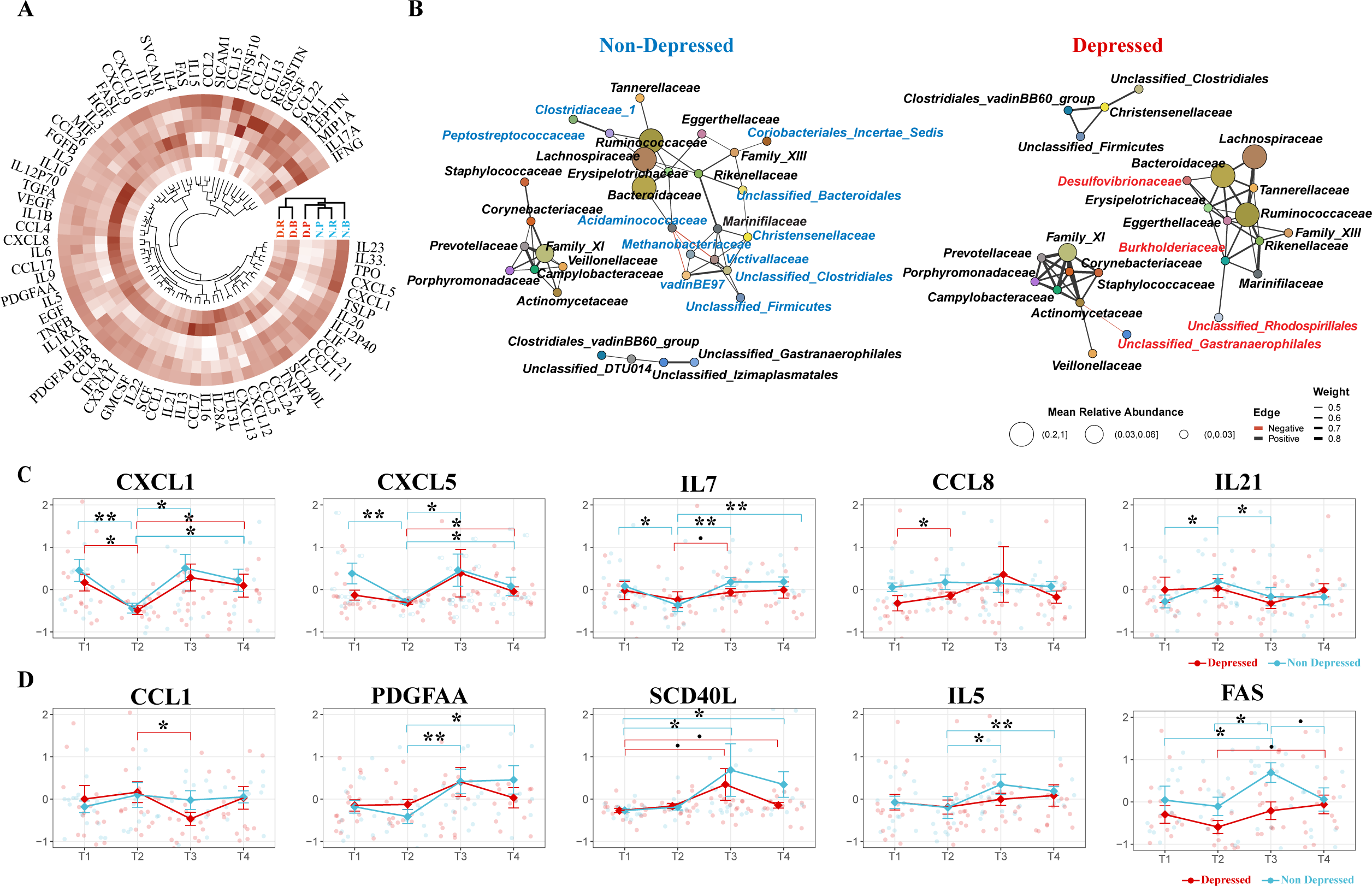
Association of Prevotella Enterotype with Reduced Inflammation and Increased Gut Microbiome Stability. **(A)** Hierarchical Clustering of Cytokine Values. Clustering based on enterotype and depression status, illustrating the cytokine profiles across different groups. **(B)** Co-occurrence Network of Microbial Families. Network representations comparing depressed and non-depressed individuals. The left network represents the non-depressed group, and the right network represents the depressed group. Unique microbial families to each network are color-coded (blue for non-depressed, red for depressed). **(C)** Immediate Changes in Cytokines Post-Intervention. Significant Cytokine Changes Between T1 and T2. Identification of cytokines that showed significant differences between T1 and T2 in at least one group (depressed or non-depressed). **(D)** Delayed Changes in Cytokines Post-Intervention Significant Cytokine Changes Between T2 and T3: Identification of cytokines that showed significant differences between T2 and T3 in at least one group (depressed or non-depressed). A pairwise two-sided Student’s T-test was used for comparing cytokine levels between different time points. Significance levels are indicated as follows: BH-adjusted p < 0.1 (.), BH-adjusted p < 0.05 (*), BH-adjusted p < 0.01 (**), BH-adjusted p < 0.005 (***).

Cytokines and chemokines, which typically spike during inflammatory states, are crucial in regulating the gut microbiome’s stability^98^, the overall different cluster on cytokine profile may signal differing microbiome ecologies between populations. Therefore, the observed differences in cytokine profiles between the two populations may reflect variations in their microbiome ecology such as stability measured by co-occurrence^102,103^. To delve deeper into this association, we constructed a family level^104^ co-regulatory network of the gut microbiome for both depressed and non-depressed groups. Mirroring our earlier result, the family of two driver genus of two major enterotype, *Bacteroidetes (family Bacteroidaceae)* and *Ruminococcus (family Ruminococcaceae)*, formed a major module network, whereas *Prevotella (family Prevotellaceae)*, typically characterized by a binary distribution among populations, formed a distinct module (**Fig. 2A, Fig. S2A, Fig. 3B**). The network analysis revealed eleven bacterial families uniquely associated with the *Bacteroidaceae-Ruminococcaceae* module in non-depressed individuals, and three distinct to depressed participants within the same cluster; however, the *Prevotellaceae* cluster showed similar patterns in both groups. The observed shifts in inter-dependency patterns within the gut microbiome between depressed and non-depressed individuals indicate variations in microbiome stability, particularly under the dominance of *Bacteroidaceae* or *Ruminococcaceae* compared to *Prevotellaceae*. Notably, such altered co-occurrence patterns of gut microbiome have been linked to individual’s diminished responses to antidepressants^73^, and our findings suggest they may also predict lower well-being scores among individuals with depression.

The interdependency of the gut microbiome between individuals initially classified as depressed and those who are not reveals interesting biological implications. In our study, core microbiome clusters in non-depressed individuals predominantly included families like *Peptostreptococcus* and *Clostridiaceae* which are known for their short-chain fatty acid (SCFA) production^105,106^, beneficial for gut health and anti-inflammatory responses. Conversely, the microbiome in depressed individuals exhibited a significant presence of opportunistic pathogens, notably *Desulfovibrionaceae*. (**Fig. 3B**). The genus *Desulfovibrio* within this family, implicated in diseases such as inflammatory bowel disease (IBD)^107^, depression^88,108^, and obesity^109^, may contribute to these conditions through its production of hydrogen sulfide^110,111^ and immunogenic lipopolysaccharides (LPS)^112,113^. These substances are known to inflict inflammation-induced damage to the blood-brain barrier^114,115^ and enhance intestinal permeability^116–118^, illustrating a possible pathway by which alterations in the gut microbiome can influence systemic inflammation and, consequently, mental health.

Despite our PERMANOVA not showing a significant variance in cytokine levels over time overall, we pursued the identification of specific cytokines with significant temporal shifts. Using a two-sided Student’s T-test, we identified five cytokines that showed significant alterations at T2 and/or T3, indicative of short and/or long-term effects associated with the program. Notably, CXCL-1 (GROA), demonstrated a consistent decrease at T2 across both groups (**Fig.3C**). CXCL-1 is noted for its mechanistic involvement of brain disorders such as Alzheimer’s disease^119,120^ and multiple sclerosis^121–123^; more importantly, it has been directly associated with the development of depression, as evidenced in both animal models^124–126^ and human clinical studies^127–129^. Following the nine-day immersive psychosocial intervention program, the cytokines CXCL-5, IL-21, IL-7, and CCL-8 exhibited immediate perturbations. In contrast, PDGF-AA, SCD-40L, IL-5, CCL-1, and FAS exhibited changes two weeks post-program, indicating a delayed response. (**Fig.3D**).

The cytokines identified in our analysis, many of which are linked to the pathogenesis of depression^130–132^, highlight the complex interplay between inflammation and mood disorders. The role of tumor necrosis factor-α (TNF-α) and its receptor superfamily, including FAS and CD40L, is well-established in the literature on depression^25,131,133,134^, reinforcing the theory that their involvement may pertain more to impaired tissue regeneration and neurogenesis rather than solely promoting inflammatory responses. The longitudinal observation of lower FAS levels among depressed individuals (two-way ANOVA F = 8.396, p-value = 0.0045) supports this notion^135^, suggesting a nuanced contribution of the TNF/TNF-receptor-superfamily to depression, possibly through impacts on neurogenesis. Intriguingly, the *Prevotella* enterotype group exhibited significantly lower levels of seven out of ten mentioned cytokines compared to the *Bacteroidetes* or *Ruminococcus* enterotypes, a pattern predominantly observed within the depressed cohort, except for CXCL1 (**Fig. S3A**). This finding points to a potential microbial influence on the cytokine environment, further complicating the association between gut microbiota, immune response, and mental health.

Our analysis, incorporating a mediation model to examine the interplay between gut microbiome composition (X), mental health outcomes (Y), and plasma cytokine levels (M), uncovered 179 significant mediating effects **(Table S2**), suggesting intricate relations under assumed^117,136,137^ causal frameworks. In this model, we pinpointed several bacterial genera potentially linked to depression-like symptoms, highlighting the top five genera with the strongest signals **(Fig. 4A)**. Specifically, we identified *Sellimonas* and *Bifidobacterium* as key bacterial genera associated with depression symptoms, corroborating their roles as biomarkers identified in comparative studies^40,138^ of microbiomes in healthy individuals and those with Major Depressive Disorder (MDD). Our findings also highlight *Erysipelatoclostridium’s* potential mediating role in exacerbating negative emotions via its effects on IL-33 (P = 0.016), Leptin (P = 0.028), and PAI-1 (P = 0.008) levels **(Fig.4A, Table S2)**. This finding is consistent with prior studies proposing *Erysipelatoclostridium* as a depression marker^139,140^ and its positive association with anorexia nervosa^141^ and radiation-induced intestinal injury^142^.

**Figure 4.**
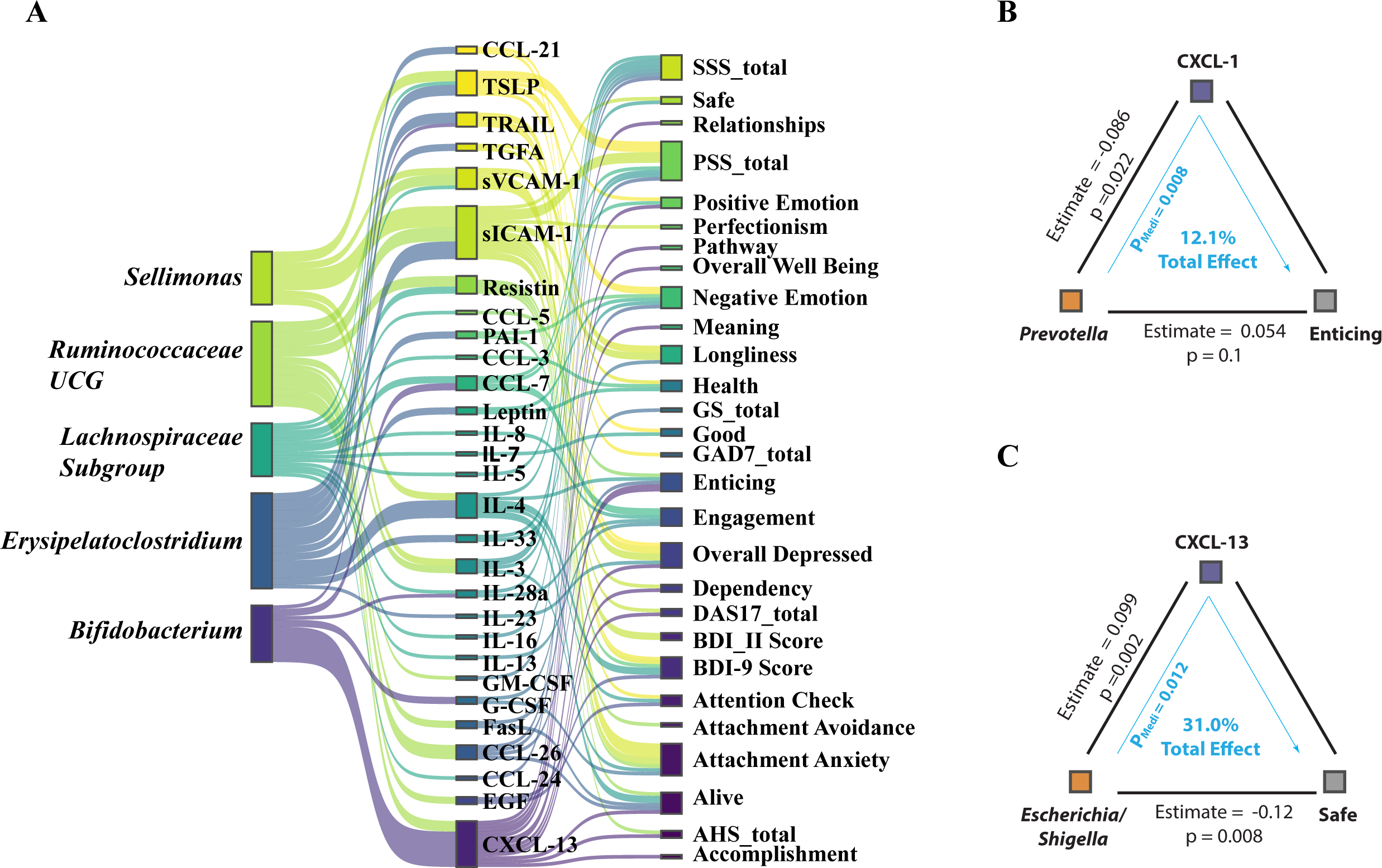
Mediation Linkage between Microbiome, Cytokine and Mental Health. **(A)** The top five bacterial genera identified through mediation analysis for their influence on mental health outcomes, mediated by cytokine levels. **(B)** The mediation association involving the psychometric parameter ’Enticing,’ the genus Prevotella, and the cytokine CXCL1. **(C)** The mediation association involving the psychometric parameter ’Safe,’ the genus Escherichia/Shigella, and the cytokine CXCL13.

These data shed light on the potentially beneficial effects of the *Prevotella* genus, notably in modulating CXCL-1 expression and its significant role in enhancing perceptions of the world as enticing **(Fig. 4B)**. CXCL-13’s mediation of feelings associated with enticement (P = 0.016) and safety (P = 0.012), and its negative correlation with the *Escherichia/Shigella* genus cluster—which is more prevalent in depressed individuals^88^—underscores complex microbiome-influenced emotional responses. **(Fig. 4B)** Further, our prior study^143^ revealed that fiber such as Arabinoxylan and LCinulin intake significantly boosts *Prevotella* levels, suggesting a dietary pathway to augment psychosocial intervention outcomes^144,145^ (**Fig. S4A**). Although these observations do not conclusively define *Prevotella* as a psych-biome marker, they open avenues for dietary interventions to potentially modulate neuro-inflammatory cytokines and manage mental stress, highlighting the intricate interplay between diet, gut microbiota, and mental health^146^.

## DISCUSSION

The pervasive link between gut microbiome dysbiosis and various mental health disorders, notably depression, has predominantly been explored through cross sectional comparisons^37,47,147^. Our investigation diverges from these traditional methodologies and represents a longitudinal examination of the microbiome’s role in mental wellness. By moving beyond simple longitudinal or cross-sectional analyses that compare depressed individuals with healthy controls, our intervention study introduces an interesting hypothesis: the existence of a *Prevotella*-dominant enterotype may contribute to a more benign inflammatory environment, specifically in relation to depression-related symptoms. This proposition is supported by our detailed examination of cytokine and chemokine profiles, including CXCL-1, which suggests a potential for mitigating inflammatory responses. Additionally, our co-occurrence analysis challenges the prevailing notion that an individual’s depression status directly influences their gut microbiome composition. Instead, we observe that the stability of the microbiome, particularly among those with *Prevotella* dominance, appears less perturbed by depressive states compared to the microbiomes of individuals with *Bacteroidetes* or *Ruminococcus* enterotypes. This distinction points to a potentially critical role of microbiome composition in moderating baseline inflammation and maintaining gut epithelial integrity under the strain of depressive conditions—a concept that underscores the importance of microbial diversity in mental health and opens new avenues for therapeutic interventions^148^.

Although the primary effect of the depression-reduction program may not be attributed to alterations in the microbiome, our findings suggest an interesting dynamic. Participants who recovered from depression did not show extensive microbiome remodeling post-intervention. In contrast, non-depressed participants exhibited more pronounced microbiome remodeling during the 9-day immersive intervention program. This finding suggests that individuals with depression at baseline may possess more static or unresponsive gut microbiomes. Such "unresponsive gut microbiomes," often characterized by low diversity, have been linked to various inflammatory conditions, including insulin resistance^98,99^, viral infections^149–151^, and cognitive decline associated with liver transplantation^152,153^. We believe that this "unresponsive gut microbiome" is a phenotype of depression-related dysbiosis, a consequence rather than a cause of depression-like symptoms.

Psychological research has long grappled with the question of whether ill-being and well-being are opposites or distinct entities. Our findings provide biological support that ill-being and well-being represent separate dimensions, each potentially influenced differently by the gut microbiome’s composition. This is particularly evident in the context of enterotypes, where *Prevotella*-dominant profiles are associated with positive emotional states^83^. *Prevotella*, known for its metabolic activity^154^, plays a role in producing neuroactive signaling molecules^115,155^, vitamins^156^, and other mood-influencing compounds^157,158^. This leads to an intriguing question: does *Prevotella* contribute to well-being through elevated production of these active signaling molecules? Our prior research^143^ demonstrated that *Prevotella* levels increase with mixed fiber intake. This finding is consistent with findings that link dietary fiber intake to a reduced risk of depression^159–161^, likely mediated by short-chain fatty acids (SCFAs) produced by fiber-digesting microbiota. SCFAs are known to regulate serotonin production^162^ and potentially other neuroregulatory molecules. This association underscores the growing interest in ’psychobiotics’ – probiotics that improve mental health through SCFA production and other mechanisms^163,164^.

Taxonomic comparisons of microbiomes between depressed and non-depressed individuals often reveal inconsistent signals^88^, attributed to the highly personalized nature of both the microbiome and mental health. Nonetheless, certain trends and mechanisms have emerged as relatively consistent across research. For example, *Coprococcus*, known for its butyrate-producing capability, consistently shows depletion in depression across numerous studies^47,147,165^, a trend confirmed by our mediation analysis. Similarly, *Sellimonas*, proposed as a depression biomarker^147^ and noted for its antibiotic resistance^166^, was highlighted in our mediation analysis. Our findings on *Bifidobacterium* also demand attention; while much psychobiome research has focused on *Lactobacillus* and *Bifidobacterium*^159,167–169^, our analysis and previous studies suggest *Bifidobacterium*’s role in psychiatric disorders may be more complex than previously thought^138,170,171^. This highlights the need for mechanistic studies to elucidate the roles of potential probiotic strains in mental health. Our mediation analysis underscores bacteria previously associated with depression, suggesting a possible link between depressive symptoms and dysbiotic gut microbiome changes.

Our findings indicate that the microbiome may influence depression through mechanisms involving immune system modulation, particularly inflammation, which has been strongly linked to depression^1,172,173^. Despite observing significant psychometric improvements in depressed individuals, their microbiome and cytokine profiles showed remarkable stability post-intervention. This persistence, even amid depression recovery, implies that these biological markers might not directly drive depressive states. Although a comprehensive analysis of inflammation’s role and its cellular underpinnings remains to be fully explored, the current data highlight subtle yet noteworthy shifts in inflammatory cytokines and chemokines following a depression reduction program, observable both immediately and over time. These alterations, though modest and not as pronounced as those seen in studies of respiratory viral infections^174^, suggest the presence of a low-grade inflammatory state rather than an acute immune reaction^175^.

Our study highlights several cytokines of particular interest in mental health research^132^. Notably, the immersive psychosocial intervention that we tested significantly reduced CXCL1 levels, a neuroinflammatory cytokine, across all participants, including those not diagnosed with depression. CXCL1, implicated in various neuropsychiatric disorders and typically elevated in depression^129,176^, has been identified as a potential therapeutic target via the CXCL1-GSK3β pathway in animal studies^124,126^. Other cytokines such as CCL17 and CXCL5 also showed reductions and are associated with depressive states in the literature^130,177^. Our mediation analysis sheds light on the role of cytokines like soluble VCAM-1 and ICAM-1 in bridging the microbiome-brain axis, crucial for maintaining the integrity of blood-brain and gut epithelial barriers^117,178^. Furthermore, cytokines including CXCL13 and IL4, known for their neuroprotective functions^179^ and ability to counter IL-1β-induced depressive-like behavior^180^, displayed significant mediative effects in our analysis. These findings highlight a nuanced role of inflammation in mental health, suggesting its potential to modulate neuroinflammatory conditions rather than exacerbating them^116^. Notably, depressed individuals with a *Prevotella*-dominant enterotype exhibited lower baseline levels of these cytokines, indicating a microbiome-immune system interaction that might favor psychosocial treatment effectiveness. This effect points to a significant interplay between specific gut microbiome compositions, immune responses, and mental health outcomes, suggesting the integration of microbiome considerations into psychosocial intervention strategies.

## LIMITATIONS

These findings should be interpreted considering several limitations. Firstly, depression was measured using self-report instruments, which, despite their common use and validity, should ideally be supplemented with clinician-rated assessments in future studies to enhance diagnostic accuracy. Secondly, the study’s scope lacks the statistical power necessary for the definitive identification of microbiome or cytokine biomarkers for mental depression. This issue is compounded by a limited sample size, which constrains the reliability of our findings, especially concerning changes—or the lack thereof—among prevalent bacterial genera. Thirdly, our mediation analysis, while offering valuable insights, rests on assumed causal links that have not been statistically verified beyond existing theoretical frameworks^136^. The potential for stress-induced microbiome alterations via cytokine pathways or the influence of other, unidentified confounding factors cannot be overlooked. Moreover, our analysis focuses on identifying potential agents of causation within the microbiome community concerning cytokine levels but stops short of categorically determining the microbiome’s impact as either beneficial or detrimental. This ambiguity stems from the personalized nature of microbiome and cytokine interactions and the absence of conclusive evidence to underpin such claims based on correlation alone. For example, the association with the Prevotella enterotype may be more indicative of dietary preferences, such as high fiber consumption, rather than a direct mood-enhancing property of Prevotella. Consequently, the associations identified herein should primarily be considered indicative of concurrent occurrences rather than direct biological mechanisms. To establish definitive mechanisms, further research employing more rigorously designed experimental studies is necessary, which falls outside the ambit of this current analysis.

Despite these limitations, our findings catalyze an intriguing hypothesis: individuals with depression might benefit from fostering a microbiome composition that is ’healthier’ in the context of depression-related inflammation, as explored in our research. This specific microbial configuration could predispose individuals to a more favorable inflammatory baseline, which, in turn, might enhance or correlate with the effectiveness of psychosocial therapeutic interventions tailored to depression. This hypothesis, novel in its suggestion that the gut microbiome may play a significant role in either augmenting or correlating with the outcomes of such interventions in humans, invites further exploration into the nuanced interplay between gut microbiome dynamics and inflammation in depression. This study also marks a pioneering step in elucidating the potential of microbiome-focused strategies to complement traditional mental health treatments, emphasizing the need for more targeted research into how microbial compositions influence depression-specific inflammatory processes and psychosocial wellbeing.

## METHODS

### Psychological Profiling

All participants were profiled at baseline, daily throughout the retreat, one-month later, and three months later (note: the six-month follow-up is ongoing). Participants completed psychometric surveys through REDCap evaluating mental health and well-being. For the characterization of social threat-related beliefs, the Dysfunctional Attitudes Scale (DAS-17), short-form, was used to assess social-threat-related beliefs. A subset of the Primal World Beliefs Index (PI-18), which measures underlying beliefs about the world (e.g. “The world is safe,” vs. “the world is dangerous”), was used to assess a broader subset of underlying beliefs and subsequent belief change. Additionally, the Beck Depression Inventory-II (BDI-II) was used to assess depression^1^, the GAD-7 was used to assess anxiety, and the Perceived Stress Scale was used to assess stress levels (PSS-10). Additional surveys included: PERMA profiler, Big Five Personality Index (BFI-10), Satisfaction with Life Survey (SLWS), Close Relationships Questionnaire (CRQ-36), Adult Hope Scale (AHS), Adverse Childhood Experiences (ACEs) Questionnaire, the Acceptance and Action Questionnaire (AAQ), and the Gratitude Survey (GS). The total BDI-II scores were interpreted following established guidelines^181^: scores ranging from 0 to 13 indicated no-to-minimal depression, 14 to 19 indicated mild depression, 20 to 28 indicated moderate depression, and 29 to 63 indicated severe depression.

### Blood Collection

Blood samples were obtained at four distinct time intervals to facilitate comprehensive biological profiling. Initial collections were performed on-site at the beginning (Day 1, T1) and conclusion (Day 9, T2) of the retreat. Subsequent collections at one-month (T3) and three-month (T4) intervals were facilitated through a collaboration with Phlebotek, utilizing their in-home mobile phlebotomy services to accommodate participants nationwide. Each session involved the collection of one 10-ml red top tube and one 10-ml lavender top tube for the separation and preservation of plasma, cells, and serum. These samples were immediately shipped overnight on dry ice to our Stanford laboratory, where they were preserved at -80°C pending further experimental analysis.

### Cytokines Luminex Assay

The cytokine assay employed a 76-plex kit (EMD Millipore H76), executed by the Human Immune Monitoring Center at Stanford University. The assay kits, sourced from EMD Millipore Corporation, Burlington, MA, were used in accordance with the manufacturer’s guidelines, with specific modifications as delineated below. The H76 kits comprise three distinct panels: Panel 1. consists of Milliplex HCYTMAG60PMX41BK, supplemented with IL-18 and IL-22 to create a 43-plex. Panel 2. incorporates Milliplex HCP2MAG62KPX23BK, with the addition of MIG/CXCL9, forming a 24-plex. Panel 3. features Milliplex HSP1MAG-63K, which is augmented with Resistin, Leptin, and HGF to yield a 9-plex. Samples were combined with antibody-coupled magnetic beads in a 96-well plate and incubated overnight at 4°C with shaking. Both cold and room-temperature incubation steps were conducted on an orbital shaker at speeds ranging from 500 to 600 rpm. The plates were then washed twice using a wash buffer in a Biotek ELx405 washer. Subsequently, a biotinylated detection antibody was added and incubated at room temperature for an hour, followed by a 30-minute incubation with streptavidin-PE, while shaking. After a final washing step, PBS was introduced into the wells, and readings were obtained using the Luminex FlexMap3D Instrument, with a lower limit of 50 beads per sample per cytokine. Custom Assay CHEX control beads (Radix Biosolutions Inc. Georgetown, Texas) were incorporated into all wells.

### Microbiome Data Analysis

Microbiome samples were collected using UBiome kits (UBiome, San Francisco, California), and sequencing was conducted by UBiome, employing 150bp paired-end sequencing. Data analysis was performed using DADA2 (version 1.20.0) in R (version 4.1.1), which offers advantages over UPARSE 8 in sequence analysis. Due to insufficient overlap between paired-end sequences, only forward reads were utilized for further processing. The forward primer selected for this analysis was GTGCCAGCMGCCGCGGTAA. Quality filtering parameters were set as follows: maximum allowable ’Ns’ set to zero, maximum expected errors set to two, truncation length at 150, and truncation quality at two. Taxonomic units were assigned using the DADA2 functions *assignTaxonomy* and *addSpecies* based on the 16S sequences that met the quality criteria. Relative abundances of these taxonomic units were determined by normalizing their respective read counts to the total reads at each time point. A rarefaction step was conducted, where reads were randomly sampled to a uniform depth of 10,000 reads per sample. As a result, three samples—X224325473, X467325054, X559299082—and 803 ASVs were excluded from subsequent richness and diversity analyses.

### Enterotype Analysis

The enterotypes of the microbiome samples were determined following previously established methods. Initially, sample counts were normalized to their relative abundance, and noise was filtered by retaining only features with a relative abundance exceeding 1%. Statistical dissimilarity between microbial communities was quantified using Jensen-Shannon divergence (JSD) and Kullback-Leibler divergence (KLD). A distance matrix was subsequently generated from these metrics. Partitioning Around Medoids (PAM) clustering was employed on the distance matrix to categorize the samples into distinct clusters. The optimal number of clusters (k) was determined by evaluating the Calinski-Harabasz index (CH index) for k values ranging from 1 to 20. A k value of 3 was selected based on the CH index and prior literature. Each cluster exhibited distinct microbial signatures: Cluster 1 was predominantly characterized by the genus *Bacteroidetes*; Cluster 2 mainly featured the family *Ruminococcaceae*; and Cluster 3 was primarily composed of the genus *Prevotella*. To validate the robustness of this clustering, silhouette width was calculated, offering a measure of the similarity between each sample and others within its respective cluster relative to other clusters. Additionally, Between-Class Analysis (BCA), informed by Principal Coordinate Analysis (PCoA) scores, was performed to visualize the separation between the established enterotypes. Finally, the samples were annotated with their respective enterotype classifications for subsequent analyses.

### Permutational Multivariate Analysis of Variance (PERMANOVA)

Two types of PERMANOVA analyses were conducted to explore the influence of various factors on the gut microbiome and cytokine profiles. The analyses were performed using the adonis2 function from the R package vegan.

Gut Microbiome PERMANOVA: The distance matrix for the gut microbiome was computed using the Bray-Curtis distance measure on the phyloseq object physeq_PERMANOVA. The PERMANOVA model was constructed to evaluate the effects of time points (Time) and depression status (depressed) on the microbial community bray curtis distance matrix. A total of 9,999 permutations were executed for this analysis.

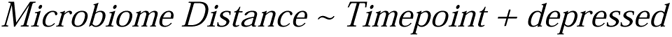

Cytokine Profile PERMANOVA: For the cytokine profiles, a distance matrix was calculated using the Euclidean distance measurements on a selected set of cytokines. The PERMANOVA model in this case was formulated to include time (Time), enterotype cluster (Enterotype), and depression status (depressed) as explanatory variables. The analysis was run with 9,999 permutations.

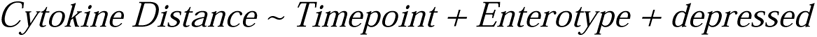

### Network Analysis of Microbiome Family Co-occurrence

To investigate the co-occurrence patterns of gut microbiome families in relation to depression, we constructed two separate networks based on individuals’ stages of depression. These analyses were performed using the R package “phylosmith” (Version 1.0.7; DOI: 10.21105/joss.01442). Initially, gut microbiome data were normalized and aggregated at the family level. This step aimed to simplify the complexity of the network by reducing the number of nodes, thereby facilitating visualization. Subsequently, pairwise Spearman correlation coefficients (rho) were computed to assess the strength and direction of associations between different microbial families. To determine an effective cutoff for rho that signifies meaningful associations in our data, we generated a null distribution of rho values through 10,000 permutations, following the guidelines provided by the authors of phylosmith. Based on this analysis, we identified cutoff values at the extreme tails (0.0001 significance level), which corresponded to rho values of 0.49 (positive association) and -0.42 (negative association). Further examination revealed that the maximum Benjamini-Hochberg (BH) adjusted p-value corresponding to these rho cutoffs was 0.00113. Given this finding, we opted for this more stringent criterion (P ≤ 0.00113) to define significant associations in our subsequent network analysis. The co-occurrence networks were constructed and visualized using the default settings in “phylosmith”. These networks illustrate the patterns of microbial family co-occurrence, providing insights into the microbial interactions that may be associated with different stages of depression.

### Two-part Zero-inflated Beta Regression Model with Random Effects

To investigate taxonomic differences between individuals classified as depressed and non-depressed in our longitudinal study, a two-part Zero-Inflated Beta Regression model with random effects (ZIBR) was utilized. Initially, data were categorized into three temporal groups: pre-treatment (T1), immediate post-treatment (T2), and extended post-treatment (T3 and T4). Data cleaning procedures involved the elimination of columns devoid of microbial counts. Additionally, taxa with fewer than four zero counts, thereby lacking a zero-inflated nature, were excluded from subsequent analyses. Following these filtering measures, the dataset was narrowed down to 20 individuals for whom data across all three temporal categories were available. This subset consisted of 11 individuals classified as depressed and 9 as non-depressed, yielding a total of 60 samples for analysis.The zibr function from the ZIBR package was employed to perform analysis on 289 distinct microbial taxa. Both the logistic and beta regression components of the ZIBR model were adjusted for depression status through the inclusion of a covariate. Hypothesis testing was conducted to assess the statistical significance of the relationship between individual microbial taxa and depression status, taking into account the zero-inflated nature of the dataset. Joint p-values were computed and subsequently adjusted using the Benjamini-Hochberg (BH) method to control for the False Discovery Rate (FDR). To further mitigate the risk of false positives, taxa appearing only once in each group (singletons) were excluded from the results.

### Mediation Analysis

Data processing was performed before running the mediation analysis to explore associations between gut microbiome (X), plasma cytokine levels (M), and psychometric parameters (Y). Specifically, only variables with a mean of non-zero values greater than 10% were retained. The gut microbiome data were then transformed using the centered log-ratio (CLR) transformation, and a prevalence filter was applied to include variables with more than 20% prevalence. Preliminary linear regression analyses were conducted to evaluate the associations between the gut microbiome and both plasma cytokine levels and psychometric parameters. Only associations meeting a P-value threshold of 0.2 were retained for subsequent analyses. Mediation analyses were then carried out using the "mediation" package in R, deploying Generalized Linear Models (GLMs) with a Gaussian distribution. A bootstrap method with 500 simulations was employed based on our previous work^136^ to estimate the Average Causal Mediation Effects (ACME) and Average Direct Effects (ADE). To assess the validity of the meditative pathways, pairs demonstrating significant ACME (P-values < 0.05) were considered to represent the indirect effects of the gut microbiome on psychometric parameters, mediated through plasma cytokine levels. Statistical comparisons between different biological conditions were also conducted to evaluate the influence of the meditative effect on each mediated pathway. Besides the traditional cutoff recommended for reporting mediation effects as the ACME P-value < 0.05, significant mediation effects were reported only when passing an additional threshold. A mediation effect was considered significant only if both the P-value for the total effect in the mediation model and the p-value from the linear model evaluating the direct effect of the gut microbiome (X) on psychometric parameters (Y) were less than 0.1. This dual-threshold method aims to add a layer of stringency to the analysis, reducing the likelihood of Type I errors. Although each test could traditionally be evaluated at P < 0.05, this conservative approach requires both to be below P < 0.1 to strike a balance between stringency and sensitivity in the analysis.

## AUTHOR CONTRIBUTION

M.P.S., G.M.S., X.Z. and A.B.G conceived the study. A.B.G, S.L. B.R coordinated the study sample collection and sequencing. X.Z. designed the overall analysis plan. X.Z., A.R., T.Y.C., H.O., performed the causal inference. J.S.J., R.H. D.J.S., performed analysis on cytokine microbiome interaction. X.L., Y.L. performed the longitudinal analysis on microbiome between groups. X.Z., G.M.S., A.B.G. and M.P.S. wrote the manuscript with the help of all authors.

## ACKNOWLEDGEMENT

We thank all participants for this study. The work is supported by NIH 5R01-MH116529, 5R25-HG010857. G. M. S. was supported by grant #OPR21101 from the California Governor’s Office of Planning and Research/California Initiative to Advance Precision Medicine. These organizations had no role in planning, writing, editing, or reviewing this article, or in deciding to submit this article for the public. X.Z. was supported by NIA fellowship Resource Centers for Minority Aging Research grant P30AG059307. We also thank support to D.J.S. from NIA-K01AG070310, and J.S.J. from the Kennedy Trust for Rheumatology Research.

Our thanks also go to Mrs. Ada Yee Ki Chen and Mrs. Lisa Stainton for their research administrative support. Our appreciation extends to the Stanford Immune Monitoring Core for their contribution to the cytokine Luminex Multiplex Assay.

We are also sincerely grateful for the generous support from Leona M. and Harry B. Helmsley Charitable Trust (grant no. G-2004-03820), the Innovative Medicines Accelerator (grant no. IMA-1051) grant, and the RAMBAM-Stanford International Collaboration grant at Stanford University.

---

**Figure S1.**
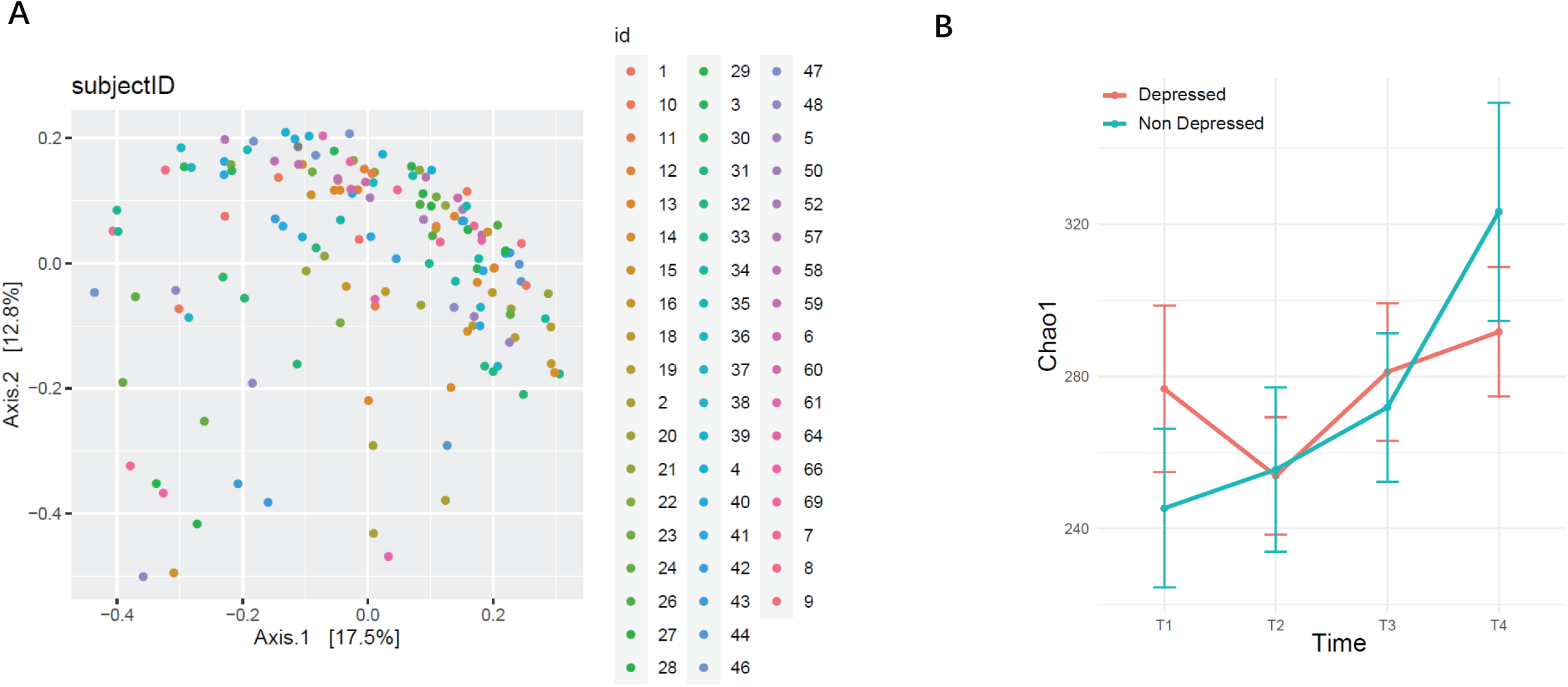
Variance of microbiome across participants. **(A)** PcoA of individual microbiome by Subject. **(B)** The Chao1 Index of depressed and non-depressed individual over time.

**Figure S2.**
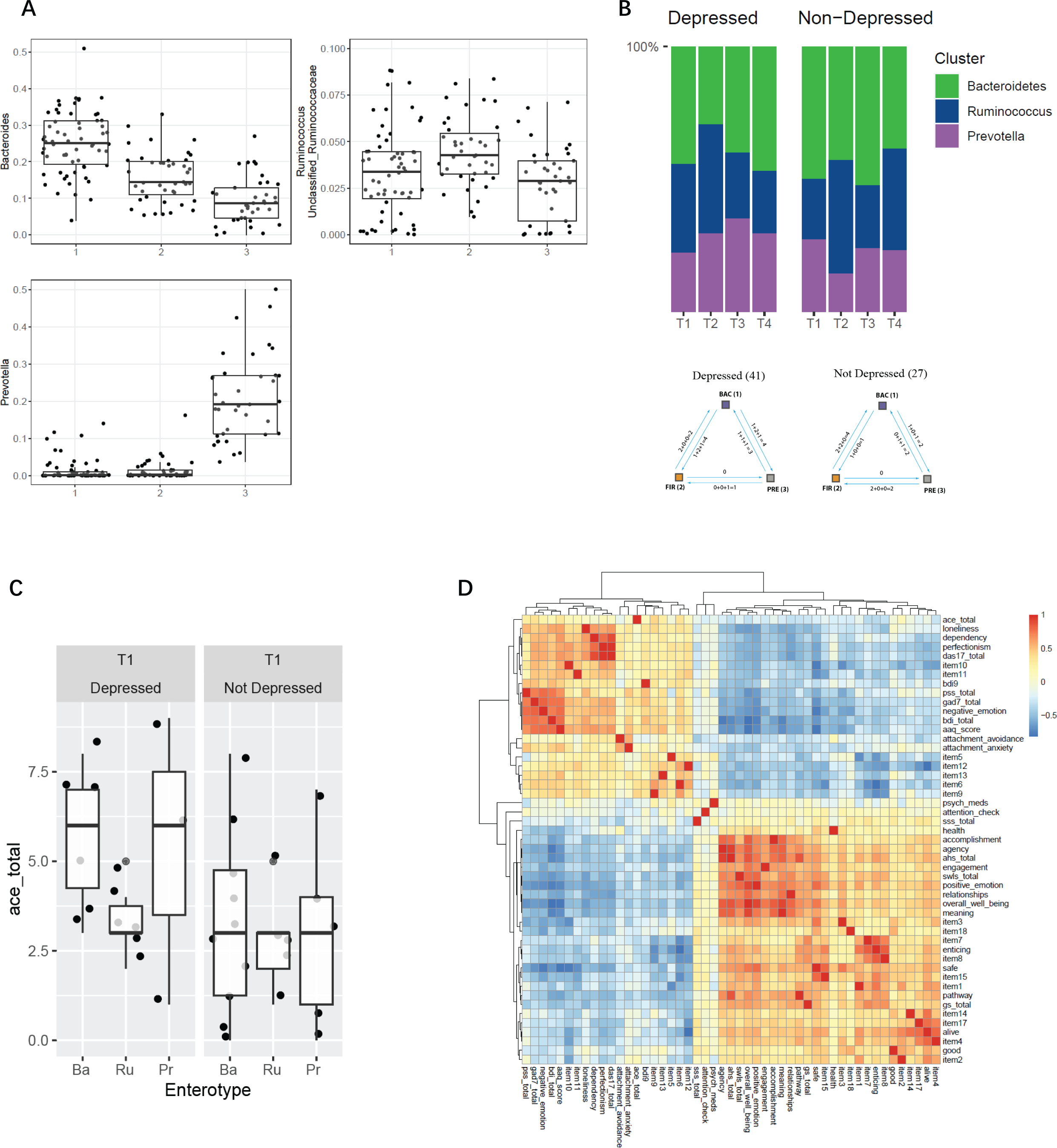
Enterotypes and its association with the program and Adverse Childhood Experiences. **(A)** The representative genera of each cluster by relative abundance. **(B)** Enterotype representation and switch dynamics through the depression group and time. **(C)** Adverse Childhood Experiences (ACE) score by enterotype and Depression status. **(D)** ACE score and other mental health measurement comparison by hierarchical clustering

**Figure S3.**
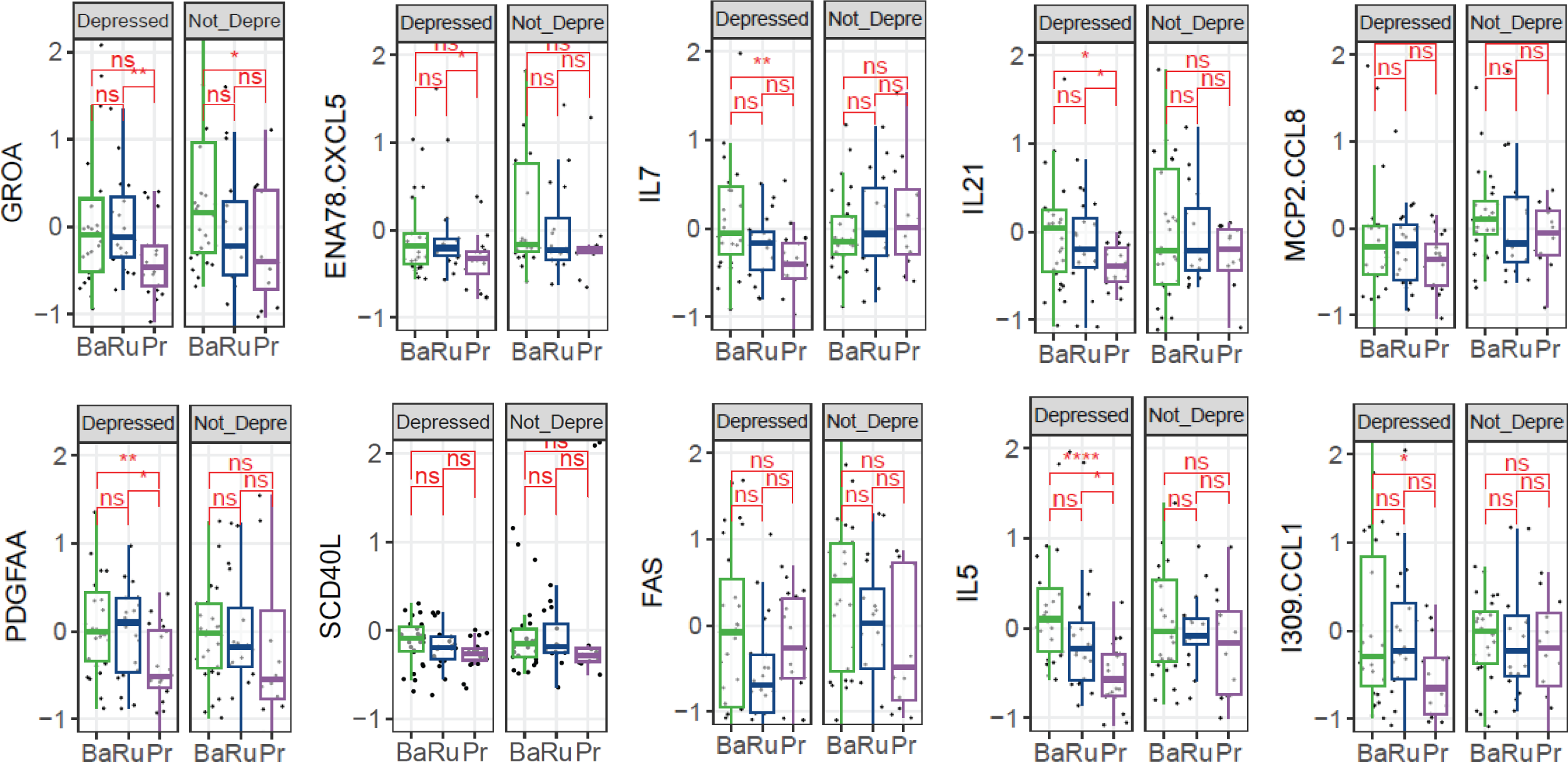
Cytokine Level Comparison Between Different Enterotype and Depression Group. The ten cytokines from Fig 3 compared across different enterotypes and between groups with and without depression.

**Figure S4.**
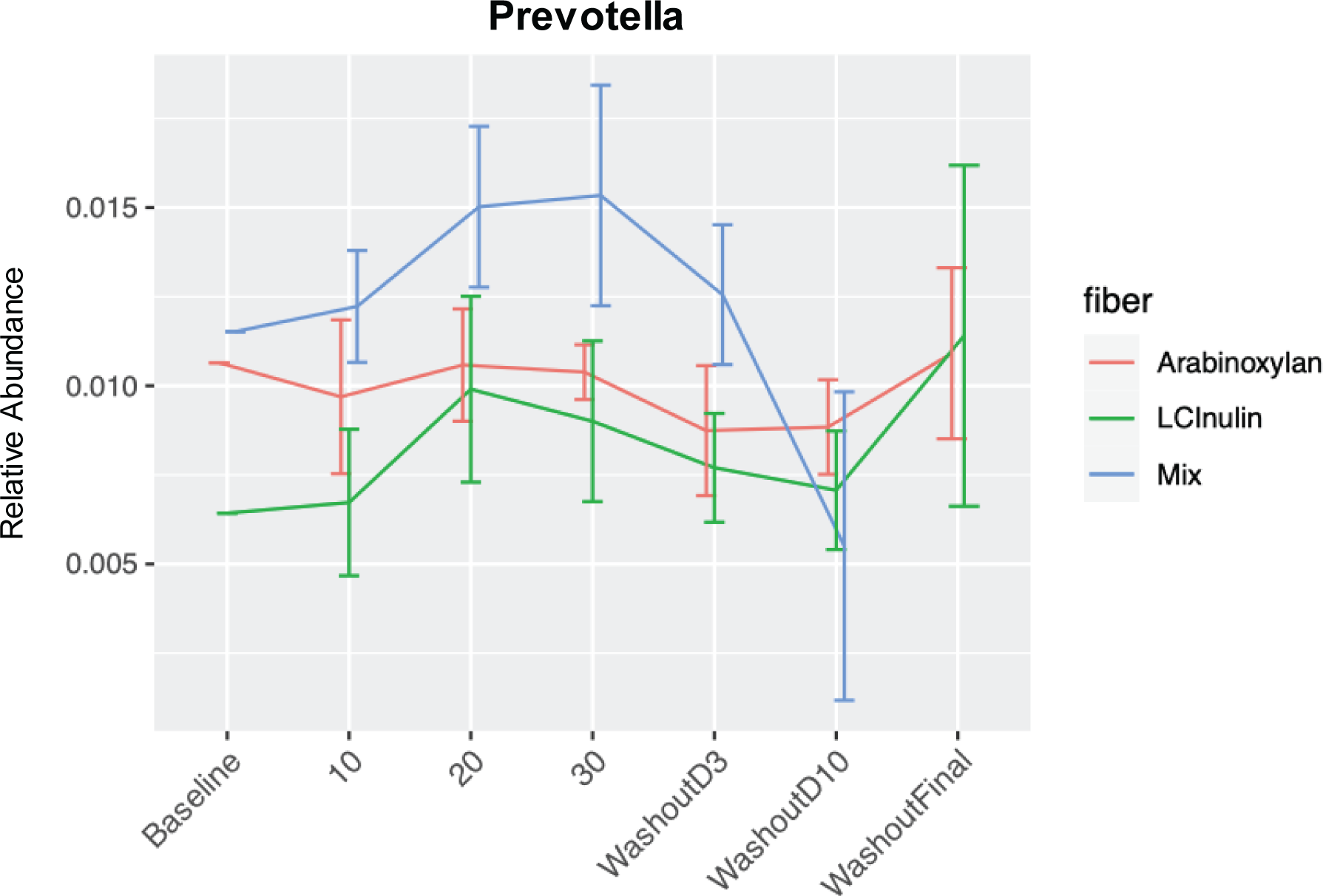
Prevotella Increase After Fiber Intake. relative abundance of the bacteria genus Prevotella in the microbiome changes after the intake of different types of dietary fiber.

